# Epigenetic clocks, sex markers, and age-class diagnostics in three harvested large mammals

**DOI:** 10.1101/2023.11.22.568330

**Authors:** Natalie Czajka, Joseph M. Northrup, Meaghan J. Jones, Aaron B.A. Shafer

## Abstract

The development of epigenetic clocks, or the DNA methylation-based inference of age, is an emerging tool for ageing in free ranging populations. In this study, we developed epigenetic clocks for three species of large mammals that are the focus of extensive management throughout their range: white-tailed deer, black bear, and mountain goat. We quantified differential DNA methylation patterns at over 30,000 cytosine-guanine sites (CpGs) from tissue samples (N=141) of all three species. We used a penalized regression model (elastic net) to build highly explanatory (black bear *r* = 0.95; white-tailed deer *r* = 0.99; mountain goat *r* = 0.97) and robust (black bear Median Absolute Error or MAE = 1.33; white-tailed deer MAE = 0.29; mountain goat MAE = 0.61) models of age (clocks). We also characterized individual CpG sites within each species that demonstrated clear differences in methylation levels between age classes and sex, which can be used to develop a suite of accessible diagnostic markers. Our results demonstrate promising tools for the large-scale estimation of age in wild animals, which have the potential to contribute to wildlife monitoring by providing easily obtainable representations of age structure in managed populations.

## Introduction

DNA methylation (DNAm) is an epigenetic modification primarily associated with the regulation of gene expression (Gallego-Bartolomé, 2020; Jung & Pfeifer, 2015). The process involves the transfer of a methyl group to or from a cytosine base (Jung & Pfeifer, 2015; Lyko, 2017; Moore et al., 2013). In mammalian genomes, this occurs primarily at cytosines that precede a guanine, also referred to as CpG sites (Moore et al., 2013). The role of DNAm in gene expression occurs at the transcriptional level where the addition of a methyl group is associated with chromatin condensation and the binding of transcriptional machinery, preventing the regular formation and activation of genes (Dhar et al., 2021; Gallego-Bartolomé, 2020; Jones et al., 2015). Likewise, the removal of a methyl group from a cytosine makes DNA available for transcriptional machinery and gene activation (Dhar et al., 2021; Unnikrishnan et al., 2019).

DNA methylation can involve predictable changes over time (Moore et al., 2013), with one such change in mammals being related to age; increasing age is associated with a global decline in DNAm (Dhar et al., 2021; Jones et al., 2015, Unnikrishnan et al., 2018). This process is thought to result from deregulation of DNAm during cell division and leads to an overall loss of DNAm as individuals age, or alternatively with the increased number of cell divisions (Jones et al., 2015; Teschendorff et al., 2013). Increasing chronological age can also be associated with site-specific increases or decreases in DNAm at predictable CpG sites within the genome (Jones et al., 2015; Unnikrishnan et al., 2019). Specific CpG sites where methylation reliably changes with age can therefore be used as biomarkers for chronological age, allowing for individual age prediction based on DNA methylation levels (Horvath & Raj, 2018; Unnikrishnan et al., 2019). The methylation-based inference of age, referred to as epigenetic clocks, is a conserved molecular mechanism across mammals (Lu et al., 2023), but species and population differences in the molecular aging process can arise due to external factors, specifically environmental conditions and life history traits (Tangili et al., 2023).

Epigenetic clocks confer many benefits when compared to traditional aging methods in mammals, such as tooth section analysis, which are typically labour intensive, invasive, and hard to implement at a large scale (Vieberg et al., 2020; Chinnadurai et al., 2016; Gasaway et al., 1978). DNAm data can be obtained from a variety of sources of DNA, including hair or fecal collection (Hao et al., 2021; Liu et al., 2021). Identifying a small number of CpG sites that strongly correlate to chronological age can facilitate the development of diagnostic markers, lowering processing costs of age estimation without sacrificing accuracy (Han et al., 2018). This approach has relevance for harvested and managed populations, where age information is often directly used to better understand processes such as survival, population growth, and harvest sustainability (Udevitz & Ballachey, 1998; Hecht, 2021; Harris & Metzgar, 1987). Free-ranging large mammals are understudied when it comes to epigenetic aging, as most non-human epigenetic aging studies focus on captive and model species (Tangili et al., 2023).

No species-specific epigenetic clock has yet been developed for the white-tailed deer (*Odocoileus virginianus*), black bear (*Ursus americanus*), or mountain goat (*Oreamnos americanus*). These three North American species hold significant cultural importance for many North American Indigenous peoples, as all three are often harvested as part of traditional food sources (Parlee et al., 2021; Schuster et al., 2012; Tryland et al., 2018). These species also contribute significantly to local economies through their involvement in hunting and associated activities. All three species are the subject of intensive management throughout their range (McShea, 2012; Hristienko & McDonald, 2007; Smith, 1988), and these efforts often rely on age information. Current aging methods for these species offer various challenges, though all three can typically be aged to at least age class by diagnostic phenotypes such as antlers and body size. Mountain goats are most often aged by the number of horn annuli, which is challenging in older animals (Stevens & Houston, 1989). Deer are typically aged by tooth wear, a method highly influenced by environmental conditions such as soil and vegetation type (Foley et al., 2021). Tooth section analysis is the preferred method for aging bears (Harshyne et al., 1998), which as noted above, is labour intensive but accurate. Further, most current aging methods require physical handling of the animal, while non-invasive aging methods, such as using body size or antler characteristics during aerial survey, are coarse and can lead to biases due to reduced detectability of certain age classes or misidentification (Davis et al., 2022). Thus, the development of epigenetic clocks could provide a significant advantage for understanding age-associated processes in these species. In this study, we developed epigenetic clocks for white-tailed deer, black bear, and mountain goat by quantifying differential DNA methylation patterns across known chronological ages. We also characterized individual cytosine-guanine sites (CpGs) within each species that were highly correlated with age class and sex to develop a suite of accessible diagnostic markers.

## Methods

### Sample Collection & DNA extraction

We collected tissue samples from three species of North American large mammals, American black bear (n=49), mountain goat (n=45), and white-tailed deer (n=47) sampled across Canada and the USA (Table 1). Mountain goat and a subset of deer samples were collected by local managers, while some deer samples in Ontario were provided voluntarily by hunters; these samples were stored immediately in ethanol. Bear tooth samples were provided by hunters via mail. All samples were then frozen upon arrival at the lab at -20°C until processing. Animals were aged to class in the field by hunters or managers, and chronological age was later estimated using either tooth section, tooth wear, horn annuli, or a combination (Supplemental Data File S1).

**Table 1.** Basic sample information, including species common and Latin names, age ranges of samples (in years), locations of sample collection, and number of individuals from each sex.

| Common Name | Latin Name | Age Range | Females | Males | Location |
| --- | --- | --- | --- | --- | --- |
| American black bear | *Ursus americanus* | 1 - 19 | 24 | 25 | Ontario, Canada |
| White-tailed deer | *Odocoileus virginianus* | 0.5 – 10.5 | 21 | 26 | Ontario, Canada;<br>Texas, USA |
| Mountain Goat | *Oreamnos americanus* | 1 – 12 | 22 | 23 | Alaska, USA;<br>Washington, USA |

DNA was extracted from samples using the QIAGEN DNeasy Blood & Tissue Kit, following the manufacturer’s standard protocol (Qiagen, Valencia, CA), and the concentration was measured using a QUBIT 3 fluorometer (Thermo Fisher Scientific). DNA was then plated in 96-well plates following an order determined using the R package Omixer version 1.6.0, which randomized plating order by covariates (i.e., age, sex) to minimize batch effects (Sinke et al., 2021). Extracted DNA was analysed using a large-scale Illumina methylation array (HorvathMammalMethylChip40) to assess DNA methylation levels at 37,492 CpG sites (Arneson et al., 2022).

### DNA methylation data and selection of species-specific CpGs

Raw DNA methylation data was provided as the intensity values for each CpG. This data was normalized and translated to beta values, defined as the ratio between methylated and unmethylated intensity, using the minfi normalization method (version 1.42.0). The ComBat function from R package sva 3.44.0, which applies parametric empirical Bayesian to account for batch effect was applied (with age as an adjustment variable). For each species the array probes were filtered to exclude CpGs that were not detected in the corresponding reference genome (black bear Accession No. ASM334442v1; white-tailed deer Accession No.

JAAVWD000000000; mountain goat Accession No. WJNR00000000). Here, species-specific CpGs were determined by aligning probes to white-tail deer, black bear, and mountain goat genomes using QuasR version 1.12.0 (Gaidatzis et al., 2015); probe sequences that did not align were discarded from subsequent analyses.

### Clock development and diagnostic CpGs

Epigenetic clock development followed the approach of Wilkinson et al. (2021). We created penalized regression models using elastic net regression within the glmnet R package (version 4.1-6) (Friedman et al., 2010). A 10-fold internal cross-validation on the training set (black bear: n=47, white-tailed deer: n=33, mountain goat: n=40) was used to determine the optimal penalty parameter (λ). In addition to fitting models to untransformed chronological age data, two different transformations were applied and models were compared to determine optimal linear fit using median absolute error: log-transformed (log[x + 1]) chronological age, and square root transformed (sqrt[x + 1]) chronological age. We performed a leave-one-out (LOO) cross validation to obtain unbiased estimates of accuracy in regard to the DNAm age estimations, we reported as estimates the correlation (*r*) between the DNAm age estimate and estimated chronological age, and median absolute error (MAE), defined as the median absolute difference between predicted DNAm age and estimated chronological age. We also predicted DNAm age using the universal pan-mammalian epigenetic clock (Lu et al., 2023); three universal clocks were applied to each species based on a subset of probes, and the difference between predicted DNAm ages and chronological age was calculated (Δ). We compared Δ values from species-specific clocks to those from the universal clock.

Lastly, the calculated optimal penalty parameter was used to generate a list of specific CpGs that strongly correlated to age class in each study species. For each of these, DNA methylation levels across samples were plotted by age class ((i) white-tailed deer: fawn (0.5), subadult (1.5 - 2.5), adult (>2.5); (ii) mountain goat: kid (1), subadult (1 - 3), adult (>3); (iii) black bear: cub (1), subadult (2-4), adult (>4); we compared mean methylation levels between pairs of classes using a Wilcoxon test, and between all three classes using a Kruskal-Wallis test. CpGs that showed significant differences in mean DNA methylation levels between age classes were selected as diagnostic CpGs (α = 0.05). To identify CpGs diagnostic of sex, epigenome-wide association studies (EWAS) were conducted using the limma package v.3.56.2 (Ritchie et al., 2015); here we used the normalized beta values of aligned probes with age and sex as fixed effects. The most significant CpG identified by the model was selected as a diagnostic, and we compared mean methylation levels between sexes using a Wilcoxon test.

## Results

### Species-specific epigenetic clocks and diagnostic CpGs

A total of 21 samples across species were excluded from clock development due to quality control metrics (Supplemental Data File S1). The independently constructed clocks included samples from the remaining 120 individuals: 47 black bear samples, 33 white-tailed deer, and 40 mountain goat samples. The chronological age of samples ranged from 0.5–10.5 years in white-tailed deer, 1-12 years in mountain goats, and 1-19 years in black bears. Of the 37,492 probes used in the methylation array, 33,751 probes were aligned to the black bear genome, 34,070 to white-tailed deer, and 31,655 to mountain goat.

The clock using the log-transformed age model yielded the highest accuracy across species, demonstrating the lowest median absolute error (Supplemental Figure 1). Based on λ, each age prediction model identified a subset of CpGs that were predictive of age in the different species: 39 CpGs in white-tailed deer (λ =0.116), 39 in mountain goat (λ =0.143), and 31 CpGs in black bear (λ =0.297; a list of all significant CpGs can be found in Supplemental Data File S2). These final log-transformed species-specific clocks were predictive of chronological age: (i) Black bear: *r*=0.95, median absolute error or MAE=1.33 years; (ii) white-tailed deer: *r*=0.99, MAE=0.29 years; (iii) mountain goat: *r*=0.97, MAE=0.61 years (Figure 1).

**Figure 1.**
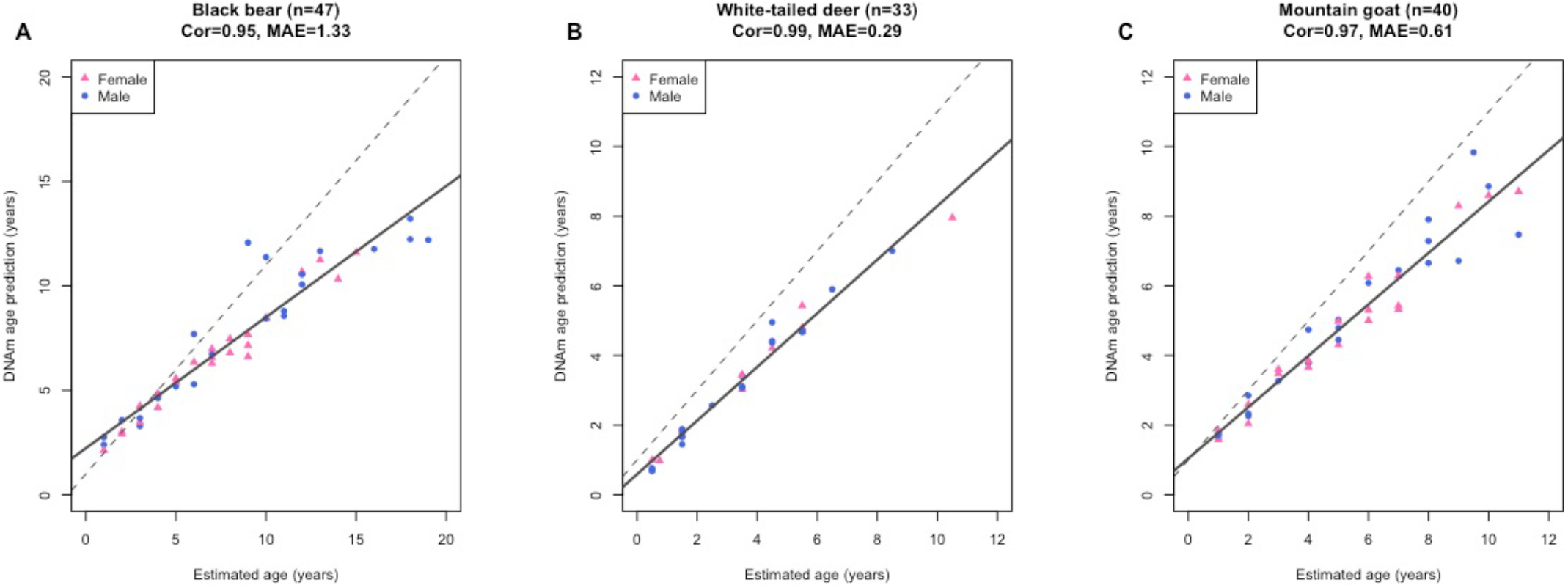
Leave-one-out cross-validation study of species-specific epigenetic clocks for **A** black bear, **B** white-tailed deer, and **C** mountain goat. DNAm age prediction (y-axis, in units of years) versus estimated chronological age (x-axis, in unit of years). The solid line indicates the linear regression of epigenetic age, and the dashed line depicts the diagonal (y = x). Cor represents the correlation coefficient (*r*), and MAE represents the median absolute error.

DNA methylation levels of the CpGs used in the species-specific clock were grouped by age class and Kruskal-Wallis and Wilcoxon statistical tests were performed (Supplemental Figure 2-4); clear differences in mean DNAm level were found between all pairs of age classes in 8 black bear CpGs, 4 mountain goat CpGs, and 2 white-tailed deer CpGs. The three CpGs in each species demonstrating the largest differences between age classes are shown (Figure 2a, c, e). Methylation levels according to sex were plotted for the most significant CpG identified by our model for each species, and a pairwise comparison between sexes was conducted using a Wilcoxon statistical test; significant differences were found between the mean methylation levels of each sex for all three species (Figure 2b, d, f)

**Figure 2.**
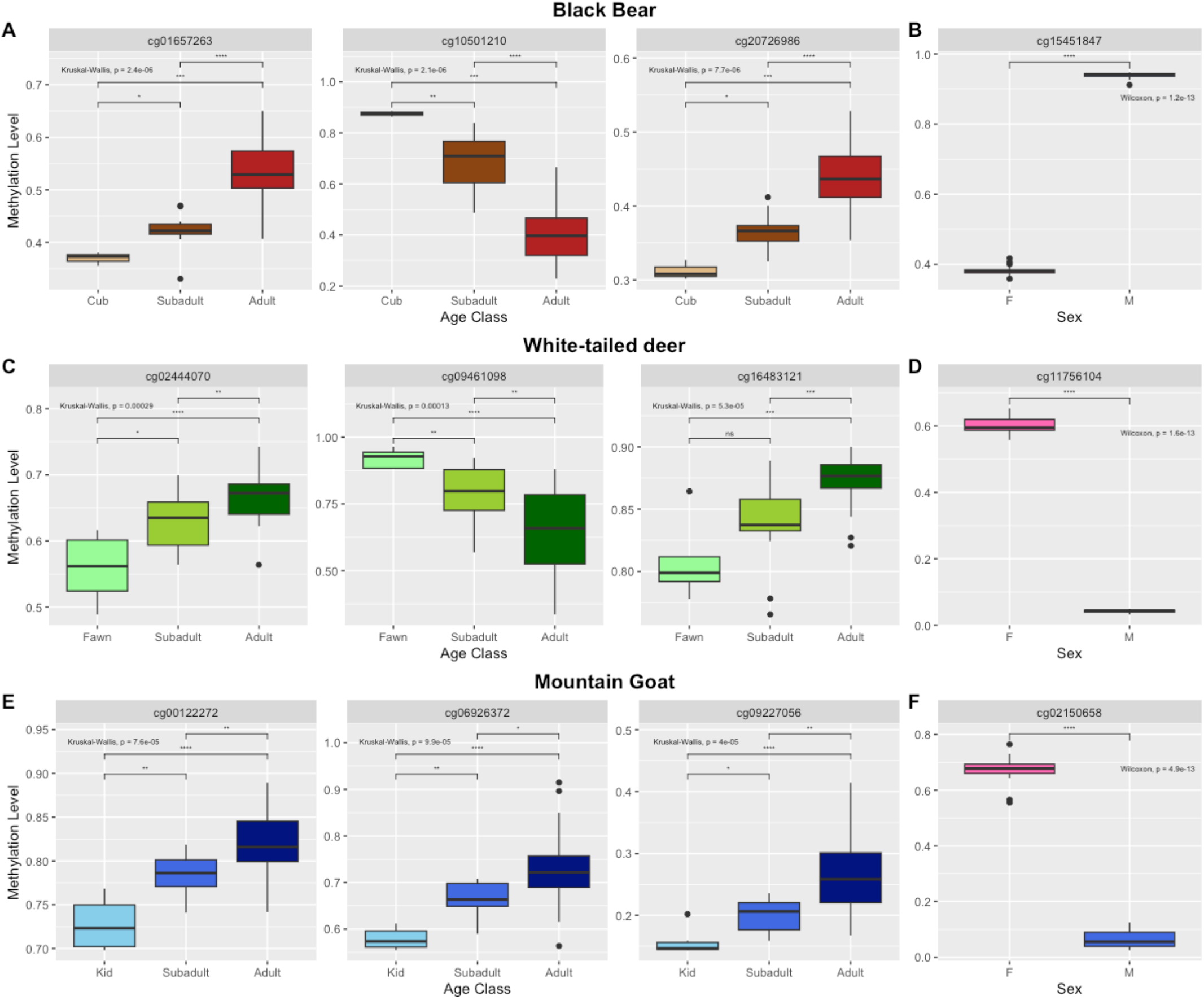
Boxplot of methylation level across age classes (**A, C, E**) and sex (**B, D, F**) at specific CpGs in black bear, white-tailed deer, and mountain goat. P-value significance level for pairwise comparisons (Wilcoxon test) is represented by asterisks (*). P-value for the comparison between all three age classes (Kruskal-Wallis test) and between sexes (Wilcoxon) is reported for each CpG.

### Universal clock comparison

We applied three previously published universal pan-mammalian clocks to each species and compared the resulting predicted age to known chronological age to determine accuracy (Lu et al., 2023). Of these three clocks Clock 1 performed poorly, reflected in high Δ values and negative age predictions (Supplemental Data File S3). Mean Δ values were lowest for Clock 2 and were as follows: 2.27 for black bear, -0.83 for white-tailed deer, and -2.66 for mountain goat (Figure 3a-c). Overall, species-specific clocks showed lower Δ values (Figure 3d-f).

**Figure 3.**
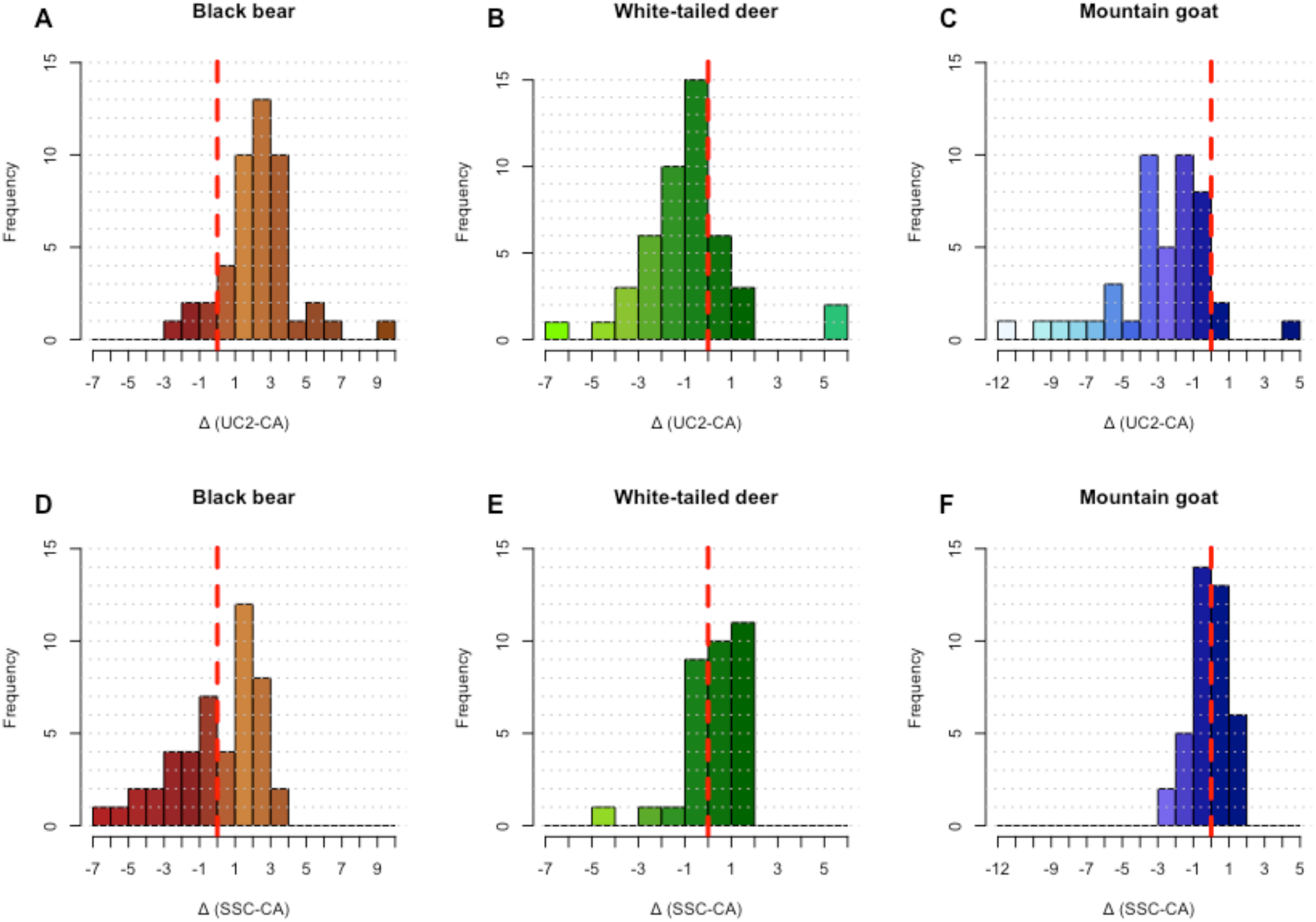
Comparison of Δ values (predicted DNAm age - chronological age) calculated from the universal pan-mammalian Clock 2 (**A – C)** and the species-specific clocks (**D – F**). The dashed red line represents a value of 0 (no difference between ages).

## Discussion

We report the development of novel species-specific clocks for three harvested large mammals in North America: white-tailed deer, black bear, and mountain goat. We also identified individual diagnostic CpGs for age-class and sex that negate the need for a genome-wide array; for example, the requirement of just a handful of CpGs to sex and identify the age-class greatly increases accessibility and decreases cost. Further, the developed species-specific clocks represent a tool for estimating the age of these three species with low error and invasiveness, while circumventing reliance on diagnostic phenotypes. Importantly, these clocks can be applied to samples that cannot be aged using traditional methods (e.g. a mountain goat with broken horns or butchered and processed animals).

The species-specific clocks demonstrated reduced error in predicted ages when compared to the published pan-mammalian epigenetic clocks, which is consistent with literature that suggests species-specific clocks improve the accuracy of age predictions (Peters et al., 2023). This finding is likely reflective of models built specific to the species, which excludes > 300 CpGs used in universal clocks including those not present in a species genome. While the clocks we developed share similar patterns to other mammals (e.g. Caulton et al., 2021; Robeck et al., 2023), the residuals appear to increase with age, notably in bears and mountain goats, suggesting reduced accuracy in older individuals. This is possibly due to error in the chronological ages used in our model. Age-related decline in the accuracy of traditional aging methods is common in all three study species (Stevens & Houston, 1989; Harshyne et al., 1998; Storm et al., 2014; Foley et al., 2021). External factors like disease (e.g., Bobak et al., 2022), inbreeding (Larison et al., 2021) and stress (Zannas et al., 2015; Pacht et al., 2021) can also influence biological aging in the form of DNAm and could also be contributing uncertainty to the model.

The majority of non-human epigenetic aging studies have focused on captive model species, leading to a gap in knowledge of wild populations of large mammals (Tangili et al., 2023). Environmental factors such as diet and stress can impact DNA methylation and aging rates (Tangili et al., 2023; Pacht et al., 2021), so the results from captive species might not be generalizable to wild populations, which experience more environmental variability (Tangili et al., 2023). In addition, large and small mammals are known to exhibit vastly different life histories and aging rates, potentially impacting the rate of DNAm (Austad, 1997; Tangili et al., 2023) and likely increasing the variance of the universal DNA methylation clock. Longer-lived species for example experience a lower rate of DNAm changes when compared to shorter-lived species, potentially due to the involvement of transcriptional regulators in epigenetic maintenance and stability (Tangili et al., 2023; Wilkinson et al., 2021). Species-specific epigenetic clocks contribute to the comprehensive understanding of DNA methylation and aging in species of various life histories, as they provide highly tailored age prediction models that more accurately represent and inform species-specific aging rates.

### A Potential New Tool for Wildlife Monitoring

Epigenetic clocks, once built, provide a reproducible and accurate tool for age prediction that has important potential for use in wildlife management. Identifying the CpGs predictive of age (<40 in our case) allows for the later development of assays that could be implemented at a relatively large scale. For example, many regions use barbed-wire hair snares for bear population estimates (Beier et al., 2010; Kendall et al., 2010); it is conceivable that with the same sample used for individual identification, a targeted DNAm assay could provide age and sex information on a population scale (e.g., Hao et al., 2021). Here, assaying a small number of CpG sites can reduce costs and equipment requirements, creating the potential for more rapid and accessible aging tools. More broadly, the use of age structure information is key in achieving one of the primary goals of wildlife management, which is to maintain harvested populations at sustainable levels. This is often accomplished by limiting or focusing harvest on specific age classes and sexes (Milner et al., 2006), which benefits from reliable data on population age structure that can be difficult or expensive to obtain. For example, the inclusion of calves and yearlings to the harvest quota of moose in Norway decreased pressure on adult females; the associated increase in average female age led to increased fecundity and population growth (Solberg et al., 1999). In other situations, the selective harvest of older individuals (i.e., those experiencing reproductive senescence) can increase the reproductive rate of a population (Milner et al., 2006). Epigenetic clocks could be used to monitor, or augment monitoring, by providing accurate estimates of population age structure which can then be used to determine if such management initiatives are warranted and to monitor their success. These clocks also may offer new avenues for developing robust estimates of age structure for species that are difficult or impossible to monitor in other ways. Ultimately, epigenetic clocks could be transformative to harvest management of cryptic species or populations at low densities that are not amenable to age classification through aerial survey. Epigenetic clock assays provide the possibility for more widespread and in-depth sampling efforts, as they rely on more easily obtainable specimens (i.e., DNA). This should lead to a more comprehensive representation of age structure within managed populations and could therefore inform more effective interventions.

## Acknowledgements

Mountain goat samples were provided by Washington Department of Fish and Wildlife (Rich Harris) and The Alaska Department of Fish and Game (Kevin White). White-tailed deer samples were collected by ABAS in Ontario and provided by Alan Cain of Texas Parks and Wildlife. Black bears samples were obtained from samples voluntarily submitted by hunters in Ontario. Species probe alignments were provided by Marie-Laurence Cossette of Trent University. This work was part of an NSERC Alliance partnership between ABAS and JN. The work was also supported by NSERC Discovery Grant (Grant Number: RGPIN-2017-03934); ComputeCanada Resources for Research Groups (Grant Number: RRG gme-665-ab); Canadian Foundation for Innovation: John R. Evans Leaders Fund and the Ontario Early Researcher Award (Grant Number: #36905). Trent University is located on the traditional territory of the Michi Saagiig Anishnaabeg; we are grateful to have had the opportunity to live and work on this land.

## Data Accessibility Statement

DNA methylation data and details on the species-specific CpGs (genome coordinates) can be found on Figshare: https://figshare.com/collections/Supporting_Files_Epigenetic_clocks_sex_markers_and_age-class_diagnostics_in_three_harvested_large_mammals/6941004

Source data underlying clock development is available in Supplementary Data Files 2-3 (S2, S3). The R software code used for this publication can be found on Gitlab: https://gitlab.com/WiDGeT_TrentU/graduate_theses/-/tree/master/Czajka/Chapter_1?ref_type=heads

## Benefit-Sharing Statement

Benefits Generated: Benefits from this research accrue from the sharing of our data and results on public databases as described above.

## Author Contributions

**Natalie Czajka**: investigation, writing-original draft, molecular laboratory work, bioinformatic analyses. **Joseph M. Northrup**: conceptualisation, sample collection, writing-review and editing, funding acquisition and supervision. **Meaghan J. Jones**: molecular laboratory work.

**Aaron B.A. Shafer**: conceptualisation, sample collection, writing-review and editing, funding acquisition and supervision. All authors edited and approved the final manuscript.

