## Supplemental Figures 1-4 for "Epigenetic clocks, sex markers, and age-class diagnostics in three harvested large mammals"

**Supplemental Data File S1.** Document containing detailed sample information, including sample ID, species names, age and age class, method by which the sample was aged, sex, tissue type, location where sample was collected, and whether the sample was included in clock development.

**Supplemental Data File S2**. Document of significant CpGs in the log-transformed species clocks, with beta values for each CpG.

**Supplemental Data File S3**. Document containing DNAm age predictions from all clocks (three universal pan-mammalian clocks, three species-specific clocks), as well as Δ calculations.

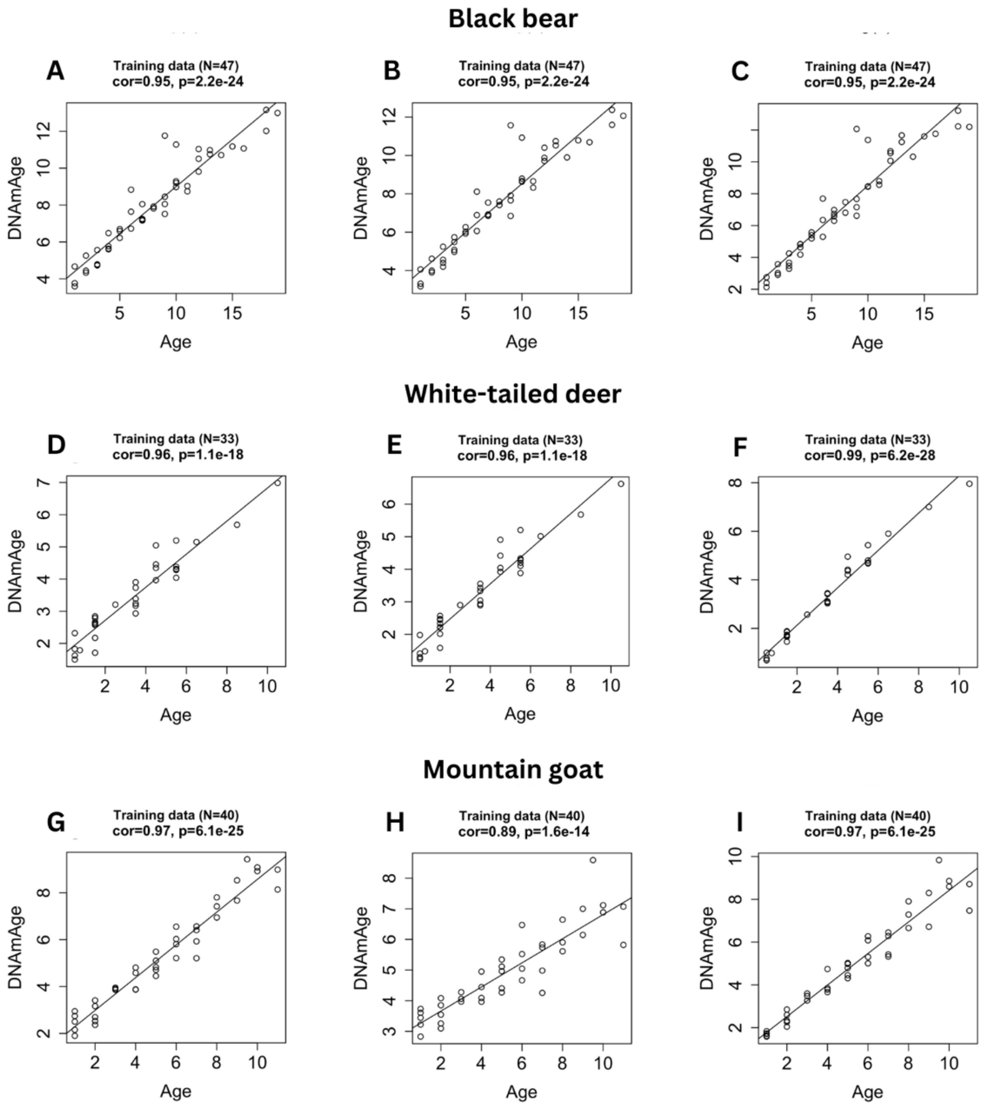


**Supplemental Figure 1.** Leave-one-out cross-validation study of species-specific epigenetic clocks for **A-C** black bear, **D-F** white-tailed deer, and **G-I** mountain goat. No age transformation **(A, D, G)**, square-root transformed chronological ages **(B, E, H),** and log-transformed chronological ages (**C, F, I**). DNAm age prediction (y-axis, in units of years) versus estimated chronological age (x-axis, in unit of years). The solid line indicates the linear regression of epigenetic age, and the dashed line depicts the diagonal (y = x). Cor represents the correlation coefficient (*r*), and "p" reports the calculated p-value.

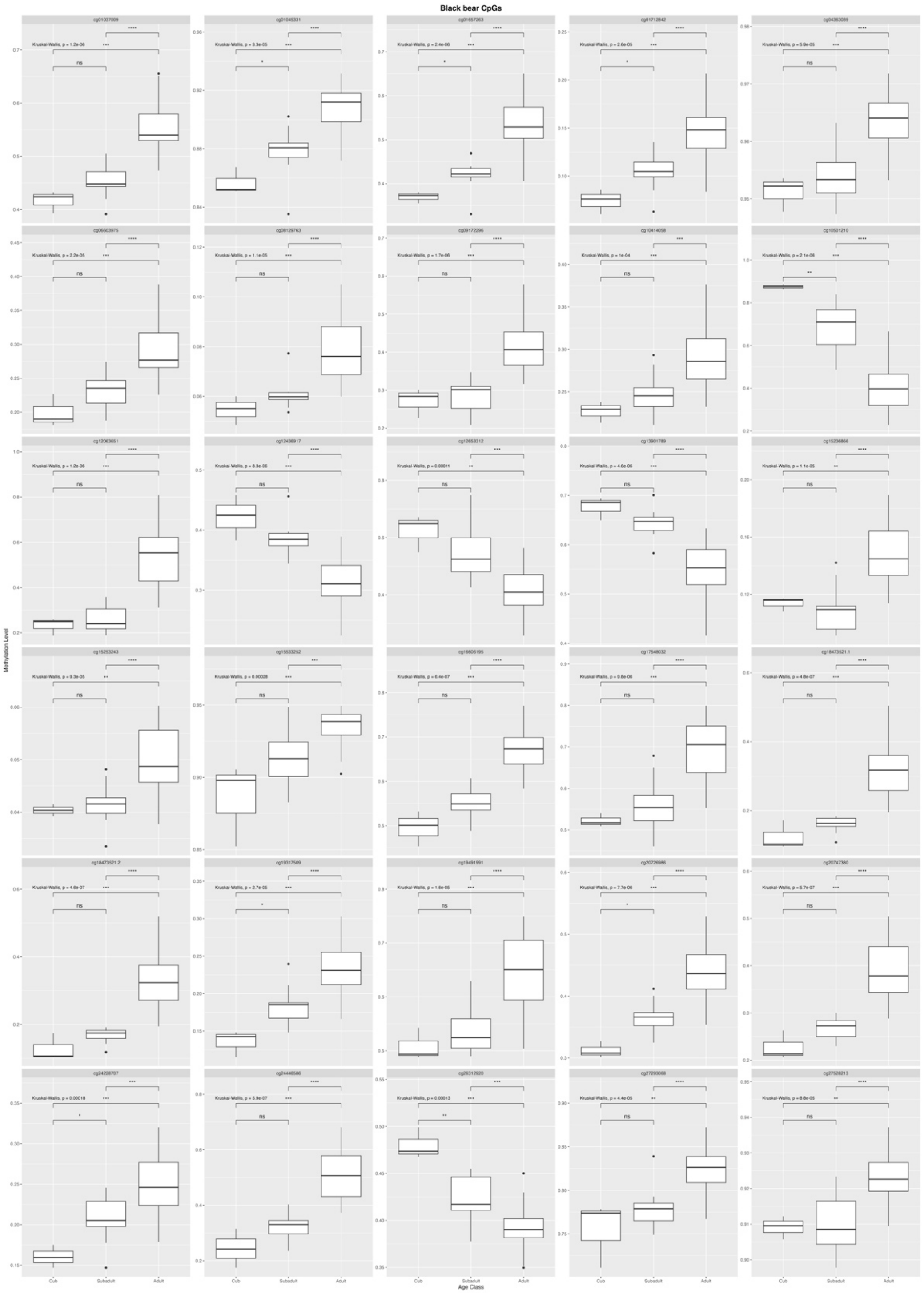


**Supplemental Figure 2.** Boxplot of methylation level across age classes at all significant CpGs in black bear. P-value significance level for the comparison between pairs of classes (Wilcoxon test) is represented by asterisks (*). P-value for the comparison between all three classes (Kruskal-Wallis test) is reported for each CpG.

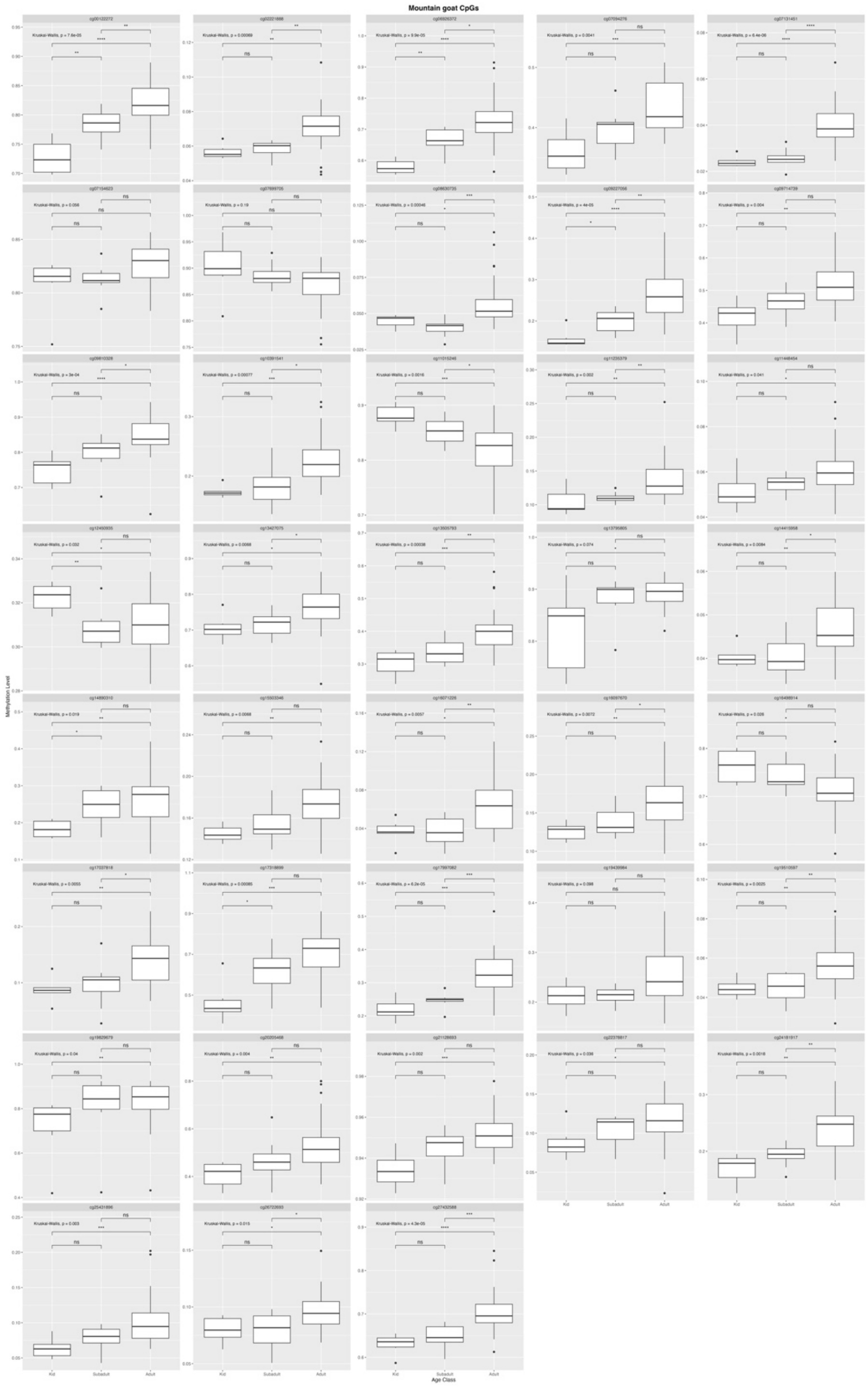


**Supplemental Figure 3.** Boxplot of methylation level across age classes at all significant CpGs in mountain goat. P-value significance level for the comparison between pairs of classes (Wilcoxon test) is represented by asterisks (*). P-value for the comparison between all three classes (Kruskal-Wallis test) is reported for each CpG.

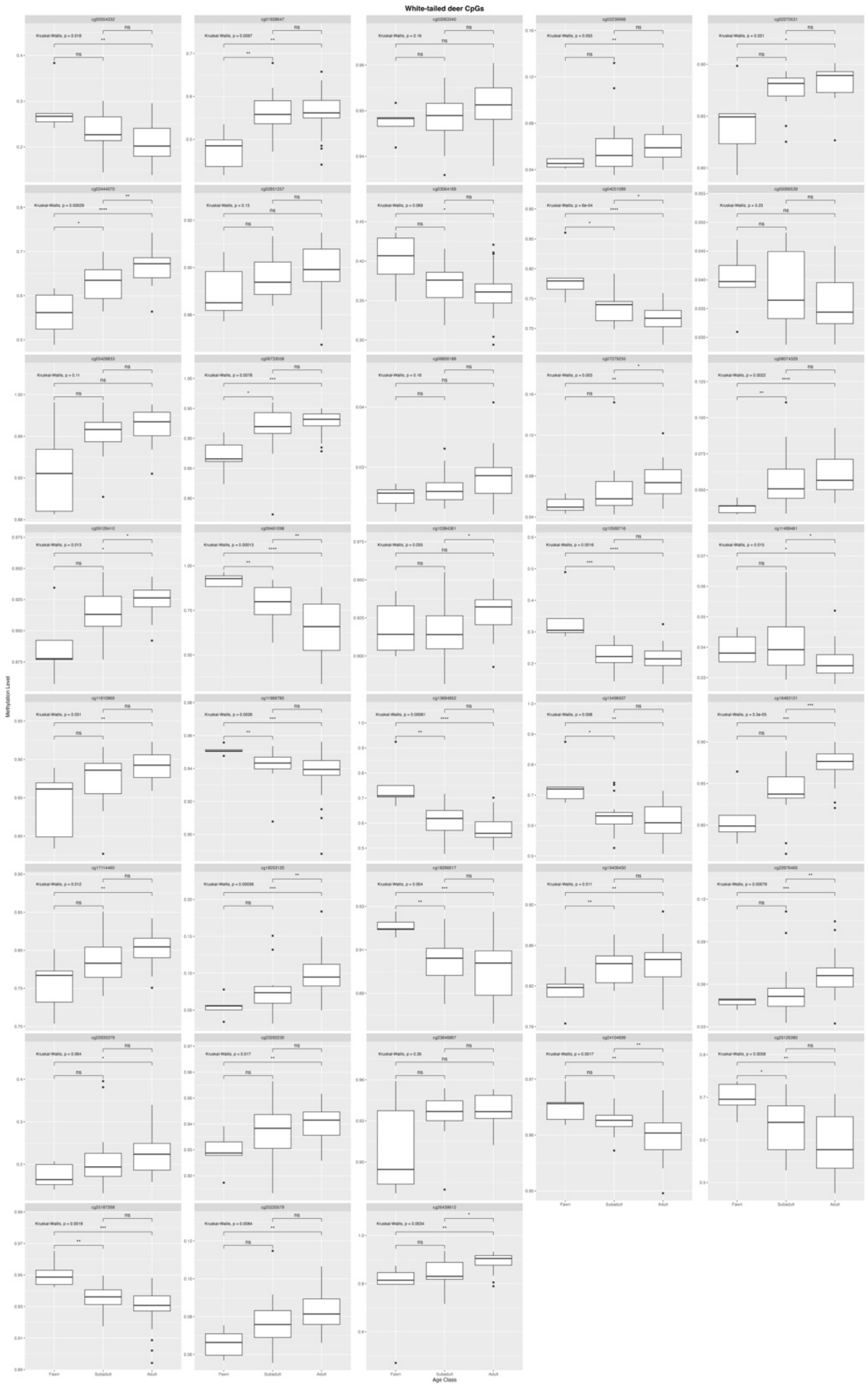


**Supplemental Figure 4.** Boxplot of methylation level across age classes at all significant CpGs in white-tailed deer. P-value significance level for the comparison between pairs of classes (Wilcoxon test) is represented by asterisks (*). P-value for the comparison between all three classes (Kruskal-Wallis test) is reported for each CpG.
